# Genome-scale profiling reveals higher proportions of phylogenetic signal in non-coding data

**DOI:** 10.1101/712646

**Authors:** Robert Literman, Rachel S. Schwartz

**Affiliations:** University of Rhode Island

## Abstract

Accurate estimates of species relationships are integral to our understanding of evolution, yet many relationships remain controversial despite whole-genome sequence data. These controversies are due in part to complex patterns of phylogenetic and non-phylogenetic signal coming from regions of the genome experiencing distinct evolutionary forces, which can be difficult to disentangle. Here we profile the amounts and proportions of phylogenetic and non-phylogenetic signal derived from loci spread across mammalian genomes. We identified orthologous sequences from primates, rodents, and pecora, annotated sites as one or more of nine locus types (e.g. coding, intronic, intergenic), and profiled the phylogenetic information contained within locus types across evolutionary timescales associated with each clade. In all cases, non-coding loci provided more overall signal and a higher proportion of phylogenetic signal compared to coding loci. This suggests potential benefits of shifting away from primarily targeting genes or coding regions for phylogenetic studies, particularly in this era of accessible whole genome sequence data. In contrast to long-held assumptions about the phylogenetic utility of more variable genomic regions, most locus types provided relatively consistent phylogenetic information across timescales, although we find evidence that coding and intronic regions may, respectively and to a limited degree, inform disproportionately about older and younger splits. As part of this work we also validate the SISRS pipeline as an annotation-free ortholog discovery pipeline capable of identifying millions of phylogenetically informative sites directly from raw sequencing reads.

## Introduction

Accurate estimates of species relationships are integral to our understanding of the evolution of traits and lineages, from modeling the co-evolution of hosts and pathogens to conserving biodiversity (Buerki et al. 2015; Bentley 2016). More accessible next-generation sequencing (NGS) technology is currently facilitating a discipline-wide shift away from resolving phylogenies using small sets of markers (typically genic/coding loci), and towards the analysis of thousands of loci from across the genome. When comparing orthologs among species, sites are said to carry phylogenetic signal when the sequence variation reflects a genetic synapomorphy that separates species or clades in a manner that reflects their evolutionary history. Conversely, non-phylogenetic signal describes sequence variation that does not reflect underlying species relationships, but rather some other phenomenon, as in cases of convergent evolution, homoplasy, or incomplete lineage sorting (Song et al. 2012; Rokas and Carroll 2008; Li, Gojobori, and Nei 1981). There was early optimism that increasing the size of datasets and selecting markers from more diverse loci would lead to the swift resolution of some of the biggest and most problematic questions in evolutionary biology (Gee 2003); yet, because non-phylogenetic signal scales up in parallel with phylogenetic signal, many species and clade relationships remain unresolved despite genome-scale efforts (Reddy et al. 2017; Philippe et al. 2011). Closer inspection of the genomic sources of phylogenetic and non-phylogenetic signal is therefore required to resolve relationships that have remained recalcitrant in the face of whole-genome data.

For decades, studies of deep-time splits have typically used more slowly-evolving protein-coding sequences (CDS), or other constrained genomic subsets, as a way to limit the impact of overwriting mutations (i.e. homoplasy) over longer evolutionary timescales (Managadze et al. 2011; Goldman and Yang 1994; Hughes and Yeager 1998; Bromham, Rambaut, and Harvey 1996; Ren et al. 2016; Salichos and Rokas 2013; Dornburg et al. 2014). In parallel, attempts to resolve more recent splits (e.g. those within a genus or family) have used sequence data from stereotypically fast-evolving loci such as intronic or intergenic regions, as these loci are thought to accrue substitutions rapidly enough to generate sufficient phylogenetic signal at shorter timescales (Debry and Seshadri 2001; Omland 2003; Baldwin and Markos 1998; Shaw et al. 2005). Studies of phylogenetic informativeness (PI) have assessed the suitability of such loci to resolve relationships (Dornburg, Su, and Townsend 2019; Su and Townsend 2015; Townsend 2007; Townsend, Su, and Tekle 2012; Graybeal 1994), often using some aspect of substitution rate as the primary predictor of ultimate phylogenetic utility (Townsend 2007; Dornburg, Su, and Townsend 2019; Klopfstein, Massingham, and Goldman 2017). These studies have found that while the relationship between rate, genomic subset, and phylogenetic utility is complicated by interacting factors including complex patterns and constraints in character evolution, tree structure, and taxon sampling (Aguileta et al. 2008; Townsend and Leuenberger 2011; Dornburg, Su, and Townsend 2019; Steel and Leuenberger 2017; Heath et al. 2008; Su and Townsend 2015), some slowly-evolving loci do appear to provide more phylogenetic signal and less non-phylogenetic signal for older splits, and vice versa (Townsend, López-Giráldez, and Friedman 2008; Fong and Fujita 2011).

However, our ability to generalize from PI studies has been limited, as individual loci contain limited amounts of phylogenetic signal and multiple factors affect that signal (Shen, Hittinger, and Rokas 2017; Klopfstein, Massingham, and Goldman 2017; Cao et al. 1994). Furthermore, a majority of quantitative studies of PI have focused on comparisons among CDS types or other historically popular markers, which has limited our understanding of other data types (Fong and Fujita 2011; Russo, Takezaki, and Nei 1996; Townsend, López-Giráldez, and Friedman 2008; Moeller and Townsend 2011; Graybeal 1994; Granados Mendoza et al. 2013; Small et al. 1998). In contrast, recent work has shown that the net PI of variable flanking regions of ultraconserved elements can rival that of protein-coding gene sequences (Faircloth et al. 2012; Gilbert et al. 2015), while non-coding introns can provide a more robust and consistent phylogenetic signal relative to coding loci (Chen, Liang, and Zhang 2017). In scaling up phylogenetic data, we continue to observe that data derived from loci experiencing distinct evolutionary forces generate topologies that are incompatible, yet often well-supported (Jarvis et al. 2014; Sharma et al. 2014; Rokas et al. 2003; Nosenko et al. 2013; Kumar et al. 2012). These conflicting results restrict our understanding of broader evolutionary processes and suggest that phylogenetic marker selection remains a critical consideration, even in this era of phylogenomics.

In this study, we compare the distribution of phylogenetic and non-phylogenetic signal among millions of sites spread across nine different mammalian locus types, including coding, intronic, and intergenic regions. The SISRS bioinformatics pipeline facilitates ortholog identification directly from unannotated whole-genome sequencing (WGS) reads, enabling genome-scale signal profiling even in non-model clades (Schwartz et al. 2015). Using WGS data from focal sets of primates, rodents, and pecora, we identify potentially informative sites without the need for traditional sequence alignment and associated substitution model correction. The vast majority of informative sites we identify derive from intronic, long non-coding RNA (lncRNA), and intergenic regions, highlighting the overwhelming phylogenetic value of these genomic subsets that have received historically limited focus. While the majority of sites from this study carry phylogenetic signal, CDS-derived sites carry the highest proportions of non-phylogenetic signal in each clade. This is especially notable given that coding sequences are often explicitly targeted for use in phylogenetics studies. Surprisingly, we find only limited support for the differential phylogenetic utility of fast- and slow-evolving locus types to resolve nodes of different ages, at least at the evolutionary timescale of this study. Thus, here we leverage our ability to identify millions of informative sites across the genome to recontextualize our understanding of phylogenetic utility among different genomic subsets, and how those trends may vary among clades.

## Methods

All associated scripts and relevant output can be found in the companion GitHub repository: https://github.com/BobLiterman/PhyloSignal_MS

### Raw Data Processing

Assessing the phylogenetic utility of genomic loci relies on having available data for species that are related under a robustly-supported underlying evolutionary hypothesis. To that end, we identified three mammalian clades with well-established evolutionary relationships (Fig. 1) and sufficient whole-genome sequencing (WGS) data: catarrhine primates, murid rodents, and members of the infraorder Pecora (Reis et al. 2018; Zurano et al. 2019; Steppan and Schenk 2017). For each clade, we obtained paired-end Illumina reads from the European Nucleotide Archive (Leinonen et al. 2011) for ten focal taxa and two outgroup taxa. To enable downstream ortholog annotation, each focal dataset contained one species with a well-assembled and well-annotated reference genome (Primates: *Homo sapiens*, Rodents: *Mus musculus*, Pecora: *Bos taurus*). We also ran a combined analysis with all taxa that we annotated using the *H. sapiens* reference genome. We assessed read data quality before and after trimming using FastQC v0.11.5 (S. Andrews – http://www.bioinformatics.babraham.ac.uk/projects/fastqc/), and raw reads were trimmed using BBDuk v.37.41 (B. Bushnell – sourceforge.net/projects/bbmap/).

**Figure 1:**
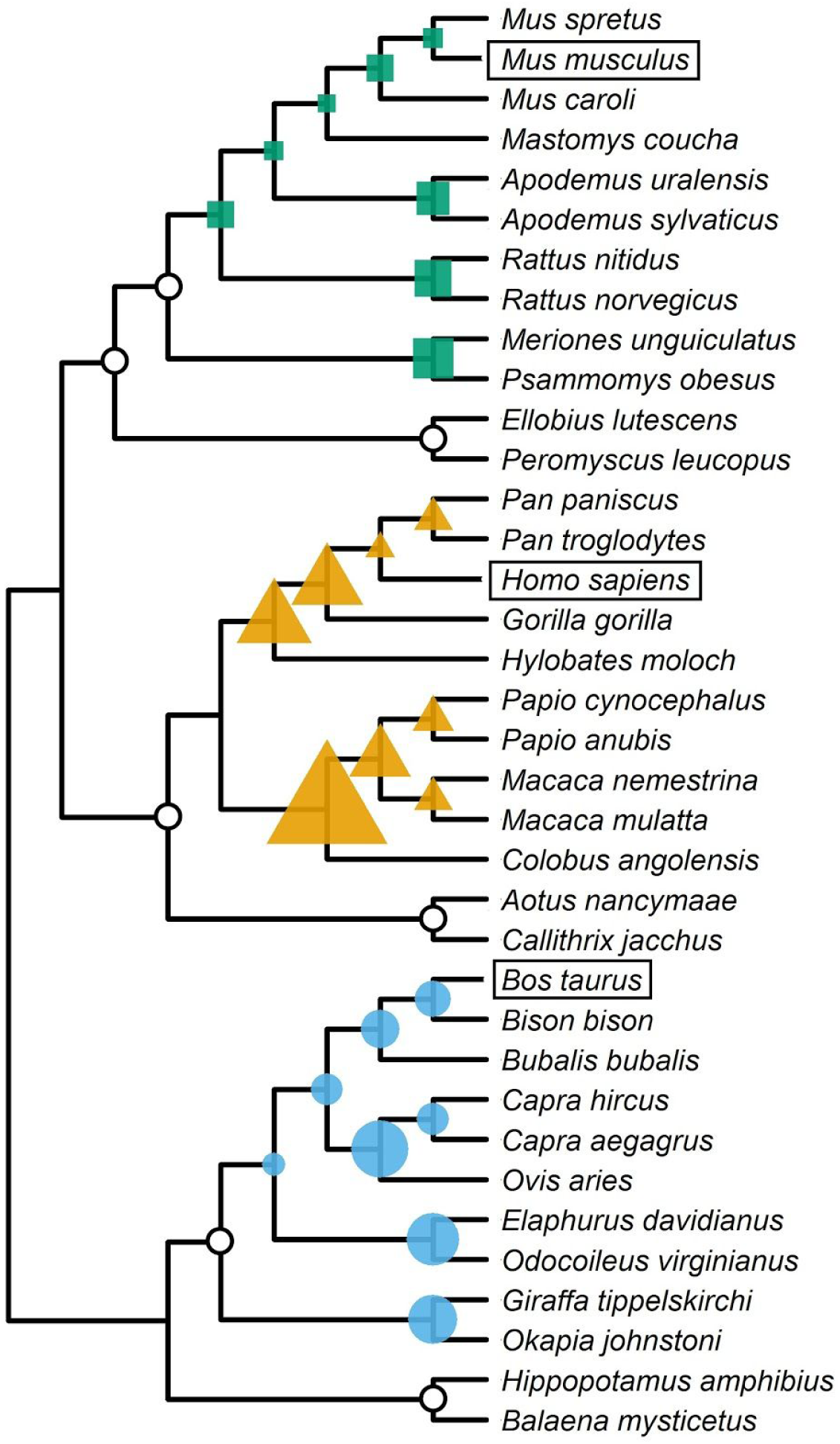
Evolutionary relationships among study species. The size of filled node icons is scaled proportionally to the number of sites supporting that split. Split support ranges for each focal group were as follows: Pecora (green squares): 172K – 1.04M sites; Primates (orange triangles): 148K – 1.87M; Rodents (blue circles): 27.4K – 600K. Open circles denote nodes included in the combined analysis that were excluded from focal analyses, and are not scaled to support size (Combined support range: 487 – 34.0K sites). Reference annotation species are boxed.

### Extracting phylogenetic data from WGS reads

For each dataset (Primates, Rodents, Pecora, and Combined), we used the SISRS pipeline to identify putative orthologs and phylogenetically informative sites (Schwartz et al. 2015). SISRS identifies orthologous loci by assembling a “composite genome” from a subsample of reads pooled across all species. By subsampling reads prior to assembly, regions of relatively high conservation have sufficient depth for assembly while taxon-specific or poorly conserved regions will fail to assemble. With a genome size estimate of 3.5Gb per dataset (Kapusta, Suh, and Feschotte 2017), reads were subsampled equally from each taxon so that the final assembly depth was ∼10X genomic coverage (e.g. 35Gb per composite genome assembly). We used Ray v.2.3.2-devel to assemble the composite genome, with default parameters and a k-value of 31 (Boisvert, Laviolette, and Corbeil 2010). This assembly represents a conserved and “taxonomically-averaged” subset of the shared regions of the genome against which all taxa can be compared.

In order to generate species-specific ortholog sets, SISRS maps the trimmed WGS reads from each taxon against their respective composite genome. Reads that mapped to multiple composite scaffolds were removed from analysis prior to composite genome conversion. When two key conditions are met (sites must be covered by at least three reads and must not vary within the taxon), SISRS uses the mapping information from each species to replace bases in the composite genome with species-specific bases. Any sites with insufficient read coverage or within-taxon variation were denoted as ‘N’. SISRS removes sites that lack interspecies variation (invariant sites), as these provide no direct topological support when inferring phylogenetic trees. For this study, we also removed alignment columns containing any missing data (e.g. Ns) as well as sites where the variation consisted of a gap and an otherwise invariant nucleotide (i.e. indels).

Finally, SISRS partitions the final alignment into overlapping subsets containing (1) all variable sites, (2) all variable sites with singletons removed (parsimony-informative sites), and (3) parsimony-informative sites with only two alleles (biallelic sites). We used the SISRS-filtered dataset containing only gapless, biallelic sites where data was present for all taxa (‘SISRS Sites’) for subsequent phylogenetic analysis.

### Data filtering and annotation

We obtained chromosome and mitochondrial scaffolds along with associated annotation data for *Homo sapiens, Mus musculus*, and *Bos taurus* from the Ensembl Build 92 database (Zerbino et al. 2018). For each reference species, we mapped their taxon-converted composite sequences onto the reference genome using Bowtie2 v.2.3.4 (Langmead and Salzberg 2012). We removed any contigs that either did not map or mapped equally well to multiple places in the reference genome, as this obscured their evolutionary origin. We also removed individual sites that displayed overlapping coverage from independent scaffolds to avoid biasing downstream results through redundant counting or by arbitrarily favoring alleles in one contig over another.

We scored each mapped composite genome site as one or more of the following eight annotation types: coding sequence (CDS, including all annotated transcript variants), untranslated regions (both 5’- and 3’ UTRs), intronic (gene region minus CDS/UTR), long-noncoding RNA (lncRNA; none annotated in Pecora), noncoding gene (genes without annotated CDS; none annotated in Pecora), pseudogenes, or small RNAs (smRNA; miRNAs + ncRNAs + rRNAs + scRNAs + smRNAs + snoRNAs + snRNAs + tRNAs + vaultRNAs). Any reference genome position that was not annotated as one of these locus types was denoted as intergenic/unannotated. In some cases, an individual site may have multiple annotations, such as lncRNA within introns, or alternative five-prime UTR regions overlapping CDS. SISRS composite sites were annotated using the reference annotation files, the output from the reference genome mapping and the intersect function in BEDTools v.2.26 (Quinlan 2014).

### Assessing annotation-specific biases in composite genome assembly and SISRS filtering

In this study, phylogenetic signal analysis was limited to (1) sites within loci that were assembled into the composite genome, and of those, (2) sites that passed through all SISRS filtration steps. For each annotation subset, we calculated the proportion of sites from the reference genome that had been assembled as part of the composite genome. We compared the relative assembly percentages among annotation types using a two-tailed modified Z-score analysis, which is robust at detecting deviations within small sample sizes (n=9 or n=7) (Leys et al. 2013; Dobbie 1963). Based on the number of annotation subsets present in each dataset, critical Z-score values indicative of significant assembly biases were identified at a Bonferroni-corrected α = 0.05/9 = 5.56E^−3^ (Primates, Rodents, and Combined; Z_Critical_ = 2.77) or α = 0.05/7 = 7.14E^−3^ (Pecora; Z_Critical_ = 2.69).

Every site assembled into the composite genome was subjected to SISRS filtration, including the removal of hypervariable sites that vary within single taxa, invariant sites among taxa, and sites with more than two possible alleles. Locus types with exceptionally high or low site removal rates were identified using the modified Z-score analysis described above.

### Assessing the phylogenetic signal among locus types from SISRS-filtered data

Biallelic sites partition a dataset into a pair of taxonomic groups, each defined by their unique fixed allele. Based on which groups of taxa were supported, the variation at each biallelic SISRS site was scored as carrying phylogenetic or non-phylogenetic signal. We tallied support for each unique split signal found in the data, including all concordant and discordant splits. For each dataset, we assessed differences in concordant and discordant site support levels using a Wilcoxon test as implemented in R v3.6.1 (R Core Team 2017). For each annotation subset, we also calculated the proportion of sites carrying phylogenetic signal and identified statistical outliers using the modified Z-score analysis as previously described.

We built phylogenies using data from the raw SISRS output (i.e. prior to reference genome mapping) as well as from each annotation-specific dataset. We inferred all trees with a maximum-likelihood approach using a Lewis-corrected GTR+GAMMA model (Lewis 2001), concatenated data, rapid bootstrap analysis, and autoMRE stopping criteria as implemented in RAxML v.8.2.11 (Stamatakis 2014). We visually assessed trees for concordance with the reference topology in Geneious v.9.1.8 (https://www.geneious.com).

### Detecting changes in phylogenetic utility over evolutionary time among locus types

To detect changes in the relative phylogenetic utility of different genomic subsets over evolutionary time, we dated each split in the reference topologies using alignment data from SISRS orthologs. For each dataset, we sorted the assembled composite orthologs based on their final SISRS site counts. The SISRS composite genome contains many contigs that are highly conserved or even completely invariant among species, so sorting by variable SISRS sites ensured that each analyzed ortholog had variation for node dating. Using mafft v.7.310 (Katoh 2002), we aligned the 50,000 most SISRS-rich orthologs and used sequence variation among these loci to estimate branch lengths on the fixed reference topology under a GTR+GAMMA model as implemented in RAxML (Schwartz and Mueller 2010). With these branch lengths, we applied a correlated substitution rate model to estimate node ages on each reference topology in R using the *chronos* function as implemented in the package *ape* v.5.3 (Paradis and Schliep 2019). To convert relative split times into absolute divergence time estimates, we calibrated specific nodes in the reference topologies using divergence time information from the TimeTree database (Kumar et al. 2017). Focal groups were calibrated at the root node using the TimeTree divergence time confidence intervals as the minimum and maximum bound estimates. In the same way, the Combined topology was calibrated at the base of the tree, but also at the corresponding nodes from the focal group calibrations. Due to stochasticity in the split time estimation process, we inferred each node age 1000 times and used the median value in all downstream analyses.

For each split in the reference topologies, we broke down the signal support by locus type (i.e. 5% of support for ‘Split A’ comes from CDS, 30% from intergenic, etc.). We used linear models in R to assess whether the relative phylogenetic utility of different locus types changed over evolutionary time. As done previously, statistical significance was interpreted at Bonferroni-corrected α values based on the number of annotation types per dataset.

## Results

### Processing of WGS reads into annotated composite genomes

Based on a genome size estimate of 3.5Gb, post-trimming read depths ranged from 10X – 38X among species (Table S1). The SISRS composite genomes contained 2M – 6M contigs, ranging from 123bp – 18Kb in length with an average N50 value of 150bp (Table S2). Through reference genome mapping, we were able to identify and annotate 32% – 79% of these sites (103M – 422M sites per dataset, Table 1, Tables S2-3). For each dataset and among locus types, we compared the proportion of sites from each reference annotation type that had been assembled and annotated in the SISRS composite genome (Fig. 2, Table 1, Table S3). Reference loci annotated as pseudogenes were significantly less likely to be assembled into the composite genome of Rodents (Z_MOD_ = 4.72; p = 2.3E^−6^) and Pecora (Z_MOD_ = 2.75; p = 5.9E^−3^), while CDS loci were assembled with a higher probability relative to other locus types in the Combined analysis (Z_MOD_ = 4.29; p=1.8E^−5^).

**Figure 2:**
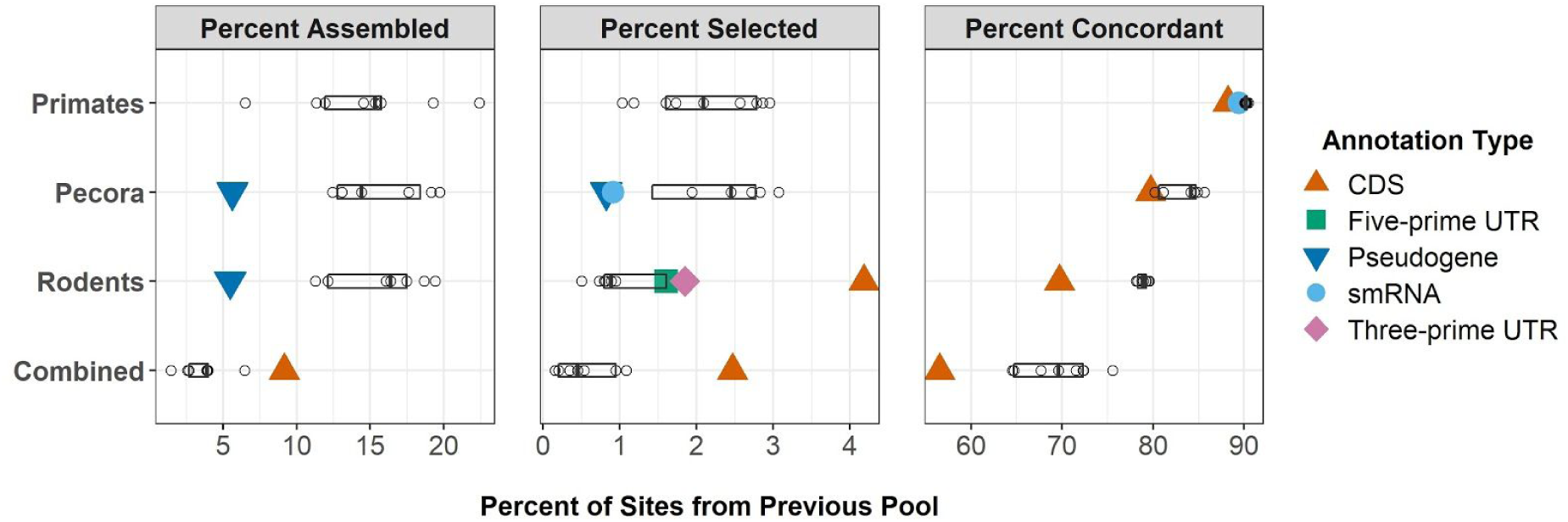
Proportion of sites passing through various steps in the SISRS pipeline. Panel 1: Based on all annotated sites in the reference genome, the percent of sites assembled into the composite genome. Panel 2: The percent of annotated, composite genome sites with parsimony-informative biallelic variation and nucleotide data for all taxa (‘SISRS sites’). Panel 3: The percent of SISRS sites carrying phylogenetic signal. Filled shapes indicate locus types with a significant deviation from the data-wide median as determined by a modified Z-score analysis. Notably, even though CDS sites were highly selected by SISRS based on their pattern of variation in rodents and in the combined dataset, CDS sites carried the lowest proportion of phylogenetic signal in all datasets.

**Table 1:** Annotation breakdown of biallelic, parsimony-informative sites (SISRS sites) and the percent carrying phylogenetic signal. Bold cells indicate annotation-specific deviations from the within-clade median, and whether the subset has significantly more (+) or fewer (-) sites than the median value is indicated in parentheses. CDS were disproportionately included by SISRS in rodents and the combined dataset, yet yielded the lowest percent concordance among all data subsets.

| Annotation | Annotated SISRS Sites |  |  |  | Percent Concordant Sites |  |  |  |
| --- | --- | --- | --- | --- | --- | --- | --- | --- |
|  | Pecora | Primates | Rodents | Combined | Pecora | Primates | Rodents | Combined |
| CDS | 156K | 94.3K | **285K (+)** | **80.6K (+)** | **79.7 % (-)** | **88.2 % (-)** | **69.7 % (-)** | **56.6 % (-)** |
| Five-Prime UTR | 5.32K | 24.2K | **19.2K (+)** | 4.40K | 85.6 % | 90.5 % | 79.2 % | 64.6 % |
| Intergenic | 6.63M | 4.47M | 1.26M | 80.4K | 84.6 % | 90.0 % | 79.5 % | 72.3 % |
| Intronic | 3.25M | 5.48M | 1.29M | 96.3K | 84.1 % | 90.1 % | 79.2 % | 71.5 % |
| Long-noncoding RNA | - | 4.12M | 886K | 128K | - | 90.2 % | 78.3 % | 67.7 % |
| Noncoding Genes | - | 18.7K | 14.1K | 763 | - | 90.4 % | 78.7 % | 72.3 % |
| Pseudogenes | **318 (-)** | 64.9K | 7.48K | 1.33K | 80.2 % | 90.3 % | 78.9 % | 69.6 % |
| small RNAs | **508 (-)** | 1.17K | 690 | 139 | 81.1 % | **89.4 % (-)** | 79.6 % | 75.5 % |
| Three-Prime UTR | 38.8K | 177K | **129K (+)** | 26.9K | 84.9 % | 90.3 % | 78.1 % | 64.72 |

### Clades vary in which locus types are selected as potentially informative (or non-informative)

SISRS called the taxon-specific base for every position in the composite genome, provided there was sufficient read support for a fixed allele (Table S4). From these sites, SISRS identified 337K – 11.8M sites per dataset with (1) sequence data for all species and (2) biallelic variation among taxa (Table S5). After reference genome mapping, we were able to annotate 300K – 11.5M of these biallelic sites (82% – 97% of SISRS sites) for use in downstream phylogenetic analyses (Table 1, Table S5). We compared the relative proportions of different annotation types that passed through the SISRS filtering process (Fig. 2, Table 1, Table S6). Among Pecora locus types, SISRS filtered out pseudogenic and smRNA sites at a higher rate than other subsets (Z_MOD_ = 3.17, 3.00; p = 1.5E^−3^, 2.7E^−3^). Conversely, CDS sites in the Combined dataset successfully passed through SISRS filtration significantly more often than other annotation types (Z_MOD_ = 8.16; p = 3.3E^−16^). The same CDS trend was even more pronounced in Rodents (Z_MOD_ = 21.1; p = 4.1E^−99^), where the five- and three-prime UTR also showed higher rates of SISRS site selection (Z_MOD_ = 4.60, 6.15; p = 4.3E^−6^, 7.6E^−10^).

### SISRS generally identifies phylogenetically useful sites

Among the three focal datasets, 78% – 90% of SISRS sites supported a split in the reference topology, and in the combined analysis, 68% of sites were concordant (Table S5). From each set of SISRS sites, concordant splits had statistically higher support than non-reference splits (all p < 2.3E^−7^, Table S7).

Phylogenies inferred for each of the three focal clades from the raw, unannotated SISRS output as well as from each locus type separately were concordant with the reference topology with high bootstrap support (all BS > 80). In the Combined analysis, the 763 noncoding gene sites were sufficient to resolve all but one node in the tree with high bootstrap support, failing to resolve the grouping of hippo and whale within the pecora. Similarly, the 139 identified smRNA sites left many relationships poorly resolved, although they did structure major groups and many sub-groups. Alignments and trees are available from the companion GitHub repository.

### Non-coding sites provide higher proportions of phylogenetic signal

Concordance percentages varied among locus types (Fig. 2, Table 1, Table S8). In each dataset we found that the CDS sites carried a significantly higher proportion of non-phylogenetic signal compared to most other locus types (Pecora: Z = 2.98, p = 2.9E^−3^; Primates: Z=11.8, p = 4.9E^−32^; Rodents: Z = 16.7, p = 2.4E^−62^; Combined: Z=4.70; p = 2.6E^−6^). Concordance percentages for CDS SISRS sites (56.6% – 88.2%) were 2.2% – 19.8% lower than the median non-coding percentage (70.6% – 90.2%) (Fig. 2, Table 1, Table S8). In addition to CDS sites, smRNA sites also carried a significantly higher proportion of non-phylogenetic signal in Primates relative to other subsets (Z = 3.00, p = 2.7E^−3^). No locus type in any dataset displayed a statistically higher percentage of site concordance (Fig. 2, Table 1, Table S8).

### Most genomic subsets show no change in phylogenetic utility over focal timescales

Estimated split times among focal groups ranged from 2.1 MY – 25.1 MY, while node ages in the Combined dataset were estimated between 1.4 MY – 46.7 MY (Table S9). We detected no significant association between relative phylogenetic informativeness and node age for seven of the nine locus types (five-prime UTR, intergenic, lncRNA, ncGenes, pseudogenes, smRNA, and three-prime UTR) (Fig. 3, Table S10). As node age increased (i.e. for older splits) the proportion of node support coming from CDS sites increased 1.4% – 2.3% per million years in the Pecora, Rodent, and Combined datasets (p = 2.46E^−3^, 3.48^−3^,2.00E^−3^ respectively, Fig. 3, Table S10). In parallel, the relative proportion of node support coming from intronic sites in Rodents and the Combined datasets decreased by 0.29% and 0.24% per million years, respectively (p = 6.04E^−4^, 2.00E^−3^, Fig. 3, Table S10).

**Figure 3:**
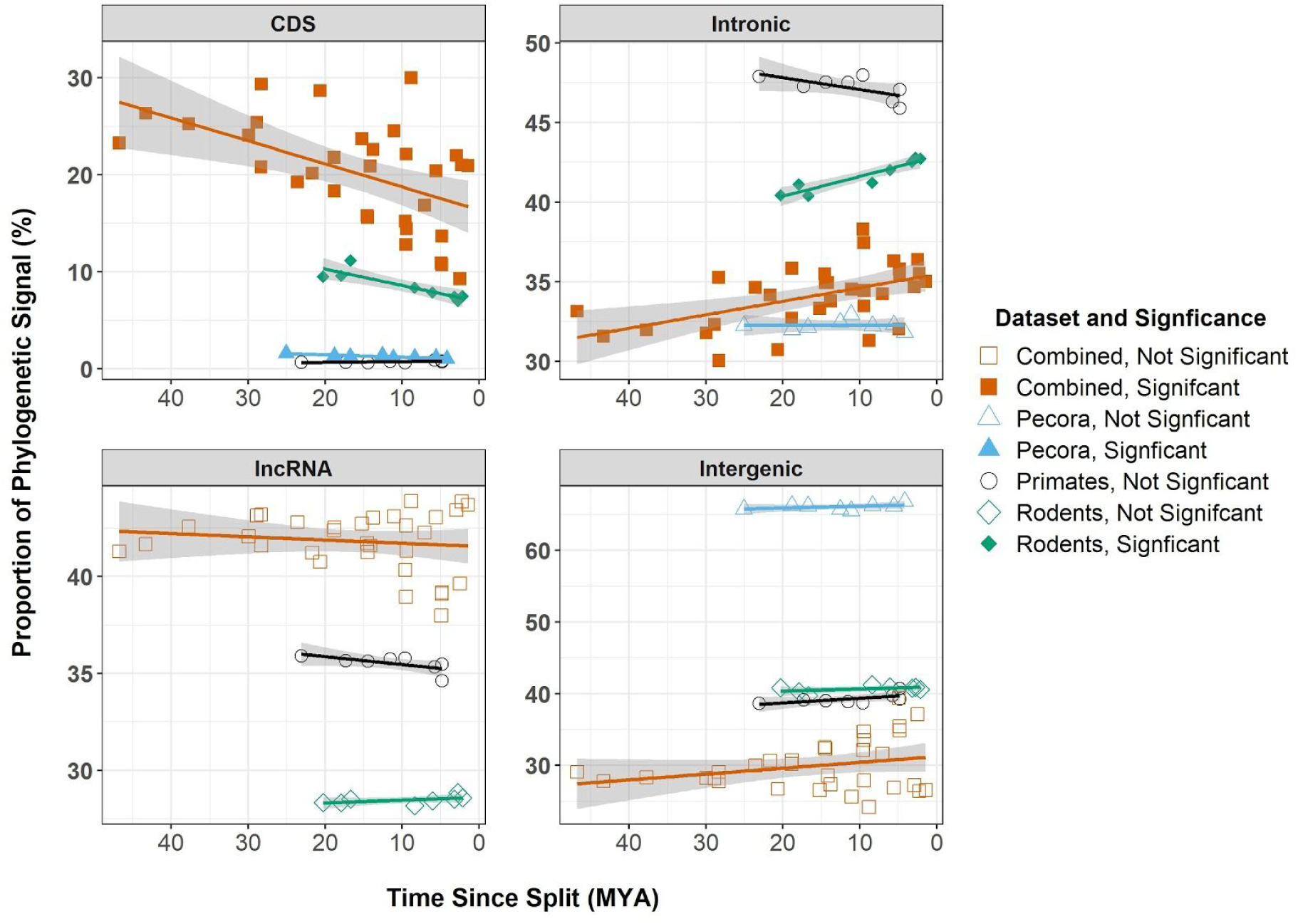
Changes in the relative phylogenetic contribution of coding sequence (CDS), intronic, long non-coding RNA (lncRNA), and intergenic sites over time. Filled shapes indicate a significant correlation between the proportion of signal coming from a locus type and the age of the node. CDS sites were more informative for older nodes in rodents, pecora, and the combined dataset, although CDS had a smaller contribution to the overall phylogenetic signal in the data. Signal from intronic regions was proportionally higher among younger nodes in rodents and the combined dataset. Sites from lncRNA and intergenic regions showed no significant change in relative utility over time. lncRNA sites are likely rolled into the intergenic fraction of pecora, as the reference genome lacks lncRNA annotation.

## Discussion

The increasing availability of publicly archived genomic data and lower costs associated with high-throughput sequencing are driving the field of phylogenetics towards genome-scale analyses; however, due in part to conflicting signal, these scaled-up datasets have not yet facilitated the resolution of many controversial relationships (Jarvis et al. 2014; Sharma et al. 2014; Rokas et al. 2003; Nosenko et al. 2013). In this study, we explore the genomic distribution of sites providing phylogenetic signal that are useful for recovering species-level phylogenies, as well as their misleading counterparts. Subsetting the data broadly by locus type (as opposed to down to the locus level, where individual loci contain limited amounts of phylogenetic signal) reduces the effect of stochastic error. Finally, analyzing only that subset of the data that displays similar patterns of variation (i.e. parsimony-informative, biallelic sites) allows us to account for and mitigate the impact of broad-sense substitution rate differences among genomic subsets; thus, facilitating equitable and straightforward comparisons of signal as a function of their genomic environment.

### Non-coding loci carry summatively and proportionally more phylogenetic signal

Coding sequences (CDS) are often used for phylogenetic analysis, due in part to the relative ease of generating sequence data (e.g. transcriptomics), combined with low mutation rates that facilitate straightforward locus annotation and alignment. However, non-coding sites in our datasets provided more total phylogenetic signal and a higher proportion of phylogenetic signal relative to CDS-derived sites in all datasets. Specifically, of the 2.4M signal-bearing sites identified in the rodent dataset, 92% were annotated as either intronic, lncRNA, or intergenic (i.e. unannotated), and this ratio rose to 98% in the larger primate and pecora datasets. Notably, given that the cow genome lacked lncRNA annotations, this high ratio in pecora suggests that the SISRS annotation-free ortholog discovery pipeline was likely able to assemble and query lncRNA loci, but that we lacked the reference information to properly identify them; hence, they were rolled into the intergenic fraction.

Given their relative abundances in the genome, that non-coding sites vastly outnumber coding sites in any truly genome-scale dataset is hardly surprising; however, CDS also carried the highest proportion of non-phylogenetic signal of any locus type in each dataset, suggesting that the same characteristics that make coding sequences an attractive target for marker development (e.g. constrained evolution) may be linked with higher relative amounts of non-phylogenetic signal (Chen, Liang, and Zhang 2017). While significantly enriched for non-phylogenetic signal in all three focal clades, CDS sites performed poorest in the rodent dataset where nearly one third of CDS sites provided support for a non-phylogenetic split, and this trend was exaggerated in the combined dataset where close to half of the CDS sites carried non-phylogenetic signal; thus, these results provide a quantitative demonstration as to why scaling up the size of datasets, especially CDS-biased datasets, may not be a cure-all for resolving phylogenetic incongruence.

### Phylogenetic informativeness of most locus types is stable across evolutionary timescales

In contrast to expectations of how markers from different genomic subsets are thought to be useful in phylogenetics studies (Dornburg, Su, and Townsend 2019; Steel and Leuenberger 2017; Phillips, Delsuc, and Penny 2004), seven of the nine locus types showed broad phylogenetic utility over the 50 million years of evolution associated with these taxa. Notably, the genomic sites expected to mutate at the most unconstrained rates (i.e. those within intergenic or pseudogenic regions) showed no significant deterioration in signal over evolutionary time in any dataset, suggesting that these subsets of the genome may provide valuable phylogenetic signal across evolutionary timescales. While the two remaining locus types (CDS and intronic regions) showed some age-dependent phylogenetic utility, these trends did not hold across all three mammal groups, even though the groups are of a similar age (20 – 25MY). For instance, CDS sites provided disproportionate support to older nodes in the pecora and rodent datasets, while there was a weaker trend in rodent introns to inform proportionally more about younger splits; yet, in primates, no locus type displayed a significant change in relative utility over time.

These contrasting results among clades may be due in part to differences in clade-specific generation times (Sims et al. 2009). Rodents, and to a lesser extent pecora, go through more generations over the same evolutionary timescale when compared to most primates. Each ‘extra’ generation provides more opportunities for new phylogenetic signal to be written, and for the existing phylogenetic signal to be overwritten. With mutations being passed down more quickly in rodents and pecora, our results concur with expectations and previous work suggesting that regions of the genome less prone to mutation (i.e. functionally-constrained CDS) may partially avoid homoplasy and provide a higher proportion of signal for older nodes in the tree. While this result may imply that CDS is of greater utility for deep-time phylogenetics as often suggested (Graybeal 1994; Ren et al. 2016; Salichos and Rokas 2013), importantly, these sites also carried the highest proportions of non-phylogenetic signal of all analyzed loci; thus, focusing on these types of markers alone could affect tree inference negatively. Together, these results suggest that decisions about which locus types provide accurate time-sensitive phylogenetic signal should be based on an understanding of biological factors that may affect how those loci evolve among focal species.

### SISRS isolates phylogenetically useful orthologs and sites directly from WGS reads

Well-assembled and annotated reference genomes currently exist for only a small fraction of life on Earth, which has limited the use of reference-guided (i.e. alignment- or mapping-based) phylogenetic marker discovery pipelines. SISRS bypasses the need for reference annotation data by assembling orthologs directly from WGS reads. In fact, millions of the signal-bearing sites identified in this study derived from completely unannotated genomic regions that would have been excluded from any reference-based ortholog discovery pipeline. Additionally, the SISRS composite genome assembly process requires only a small fraction of the per-taxon sequence data that would be required in the *de novo* development of a UCE-type dataset, making it a tractable and affordable alternative to UCEs that can be performed using previously published WGS data, as was the case for this study.

Resolving species tree discordance among datasets relies on strategies that account for or mitigate the effects of misleading non-phylogenetic signal. Here, we demonstrate that SISRS filtered data (i.e. parsimony-informative biallelic sites) is largely comprised of phylogenetically informative data, as less than a third of SISRS sites carried non-phylogenetic signal in any dataset. Notably, these sites were identified both in the absence of an underlying tree and prior to any genome annotation, suggesting the potential benefits of this framework for those working in clades that lack any substantial prior taxonomic or genomic focus. It should also be reassuring that, among datasets, 80% – 97% of SISRS sites were found in composite genome scaffolds that could be uniquely mapped back to the reference genome and annotated, as this suggests that the steps associated with SISRS filtering are effective, in parallel, at isolating primarily single-copy orthologs even in the absence of a reference genome. As genome-scale data contains nearly all the possible phylogenetic and non-phylogenetic signal, methods to address high amounts of complex non-phylogenetic signal need not rely on retaining every site for analysis.

SISRS datasets for the focal clades ranged from 3M – 11M biallelic sites, and over three-quarters of those sites provided phylogenetic signal. For the combined analysis, we identified only 300K biallelic sites; however, this dataset has no missing data and would be much larger without this limitation. Importantly, almost 70% of these sites provided accurate and useful phylogenetic signal. The reduced size of the dataset for all species (although it is still very large) may, in part, reflect the expected deterioration of phylogenetic signal over longer stretches of evolutionary time (Salichos and Rokas 2013; Pisani et al. 2012; Townsend, Su, and Tekle 2012). However, as analyses incorporate more species, the optimal rates for tree inference will shift higher as well (Townsend and Leuenberger 2011). This suggests that filtering datasets down to biallelic sites may be overly restrictive for resolving deep-time relationships. It is also important to note that the sequence diversity among all 36 species is higher than any focal subset of 12, and this likely had an impact on the composite genome assembly process, which impacts sites available to SISRS. Future refinement of the composite genome assembly process, in addition to facilitating the inclusion of tri- and possibly quadallelic sites, should allow for even more robust profiling of sequence variation from more diverse clades and/or deeper timescales.

Finally, we did not include sites with missing data in our analysis so that we could accurately characterize whether each site correctly supported the reference phylogeny. SISRS datasets grew by 50% – 100% when just a single species was allowed to be missing for any given site. Therefore, we predict that a combination of the analyses presented here (i.e. when all data is present) along with *post-hoc* analyses that utilize quartet methods will likely facilitate even more robust phylogenetic information extraction from large and complex genome-scale datasets.

## Conclusions

In this study, we provide a perspective on the phylogenetic utility of different locus types as they apply to resolving species relationships among three mammal clades. Non-coding sites provided higher proportions and amounts of phylogenetic signal compared to CDS sites in all datasets. This suggests potential benefits of shifting away from primarily targeting genes or coding regions for phylogenetic studies, particularly in this era of accessible whole-genome sequence data. Across 50MY of mammal evolution, we find that, in contrast to prior work suggesting that the phylogenetic utility of data types likely varies over time, only CDS and intronic regions varied in their utility and that these patterns were both subtle and clade-specific. As these results appear to differ from long-held beliefs about the phylogenetic utility of different genomic subsets, we hope our findings will provide the motivation to expand the search for phylogenetic markers and focus on data with the greatest accuracy for a clade, to resolve some of the more recalcitrant relationships in evolutionary biology.

